# Disrupting butterfly microbiomes does not affect host survival and development

**DOI:** 10.1101/574871

**Authors:** Kruttika Phalnikar, Krushnamegh Kunte, Deepa Agashe

## Abstract

Associations with gut microbes play a crucial role in the physiology, immune function, development, and behavior of many insects. However, butterflies may be an exception to this pattern since butterfly microbiomes do not show the host-specific and developmental shifts that are expected to evolve under strong host-microbial associations. Here, we present the first experimental test of this hypothesis by disrupting gut microbial communities of two butterfly species, *Danaus chrysippus* and *Ariadne merione*. Larvae of both the species fed on host plant leaves that were either chemically sterilized or treated with antibiotics had significantly reduced bacterial loads and disrupted gut bacterial communities substantially. However, neither host species treated this way suffered a significant fitness cost. We did not find significant variation in survival, growth and development between test larvae and control larvae. This suggested that butterflies do not rely on their gut bacteria for digestion, detoxification, resource accumulation and metamorphosis. Thus, our results provide empirical support for the growing realization that dependence on gut bacteria for growth and survival is not a universal phenomenon across insects.

## INTRODUCTION

Inter-specific interactions are crucial in shaping the ecology and evolution of organisms. This is perhaps best understood in insects, which often have specific and intimate associations with gut microbes (bacteria and fungi) that influence host biology [1–3]. The most obvious benefit provided by gut microbes is the ability to digest and survive on specific foods, potentially facilitating the use of new dietary niches. For example, termites [4,5], mosquitoes [6] and honeybees [7] rely on their gut bacteria for digestion of their typical diet. The gut bacteria of coffee bean borers [8], oriental fruit flies [9] and diamondback moths [10] detoxify the host diet, allowing survival on otherwise inedible food sources. In the western corn rootworm, gut bacteria also allow the host to make a rapid dietary shift from corn to soybean within a few generations [11]. Thus, insects have often evolved specific associations with their gut microbes that allow them to occupy a diverse range of dietary resources. It is therefore not surprising that many groups of insects have also evolved specific strategies to transmit such beneficial gut microbes across generations [12].

Butterflies present a contrast to this general pattern because they do not seem to have consistent diet-specific or stage-specific associations with gut bacterial communities. For example, multiple species of wild-caught butterflies harbor similar bacterial communities across the dramatic dietary and developmental transitions that occur during metamorphosis [13]. In addition, butterfly larvae largely mirror the bacterial communities of their diet, suggesting passive dietary acquisition of gut flora and relatively weak host-imposed selection [13,14]. Carnivorous and herbivorous larvae of lycaenid butterflies do not harbor distinct bacterial communities [15], suggesting that larvae do not depend on specific gut bacteria to consume different dietary resources. Finally, recent experimental work in the butterfly *Lycaeides melissa* showed that diet-induced variation in larval bacterial communities did not affect larval fitness [16]. These diverse studies suggest that gut bacterial communities of butterflies are mostly transient and do not have a functional association with their hosts. Here, we present a systematic experimental test of this hypothesis.

We measured the impact of gut microbes on two wild-caught butterfly species, *Danaus chrysippus* and *Ariadne merione* (figure 1). Both species belong to the family Nymphalidae and their larvae feed on toxic host plants that produce potent anti-herbivory compounds (figure 1). *Danaus chrysippus* larvae feed on a group of plants called milkweeds (family Apocynacae) [17] whereas *A. merione* larvae specialize on two plants from the family Euphorbiaceae: *Ricinus communis* (castor oil plant) and *Tragia involucrata* (Indian stringing nettle) [17]. At our study site we found that *D. chrysippus* larvae largely fed on the locally abundant milkweed, *Calotropis gigantea*. This plant produces white latex that contains cardiac glycosides – mainly Calotropin that blocks the activity of the Na+/K+ pump of herbivores [18–21]. These cardiac glycosdies are not only present in the latex, but also the foliage. Hence, the plant is poisonous to insects [18,22–24]. Similarly, *R. communis* leaves and other tissues contain the alkaloid Ricinine [25–28] that also kills insects, although the exact mode of action is unknown. Apart from such toxins, consuming plants generally poses various challenges: leaves are typically difficult to digest, and have low nitrogen content [1,29,30]. Thus, we tested whether the gut bacteria of *D. chrysippus* and *A.* m*erione* aid in digestion and dietary detoxification, as observed in other insects that feed on toxic food sources [31].

**Figure 1.**
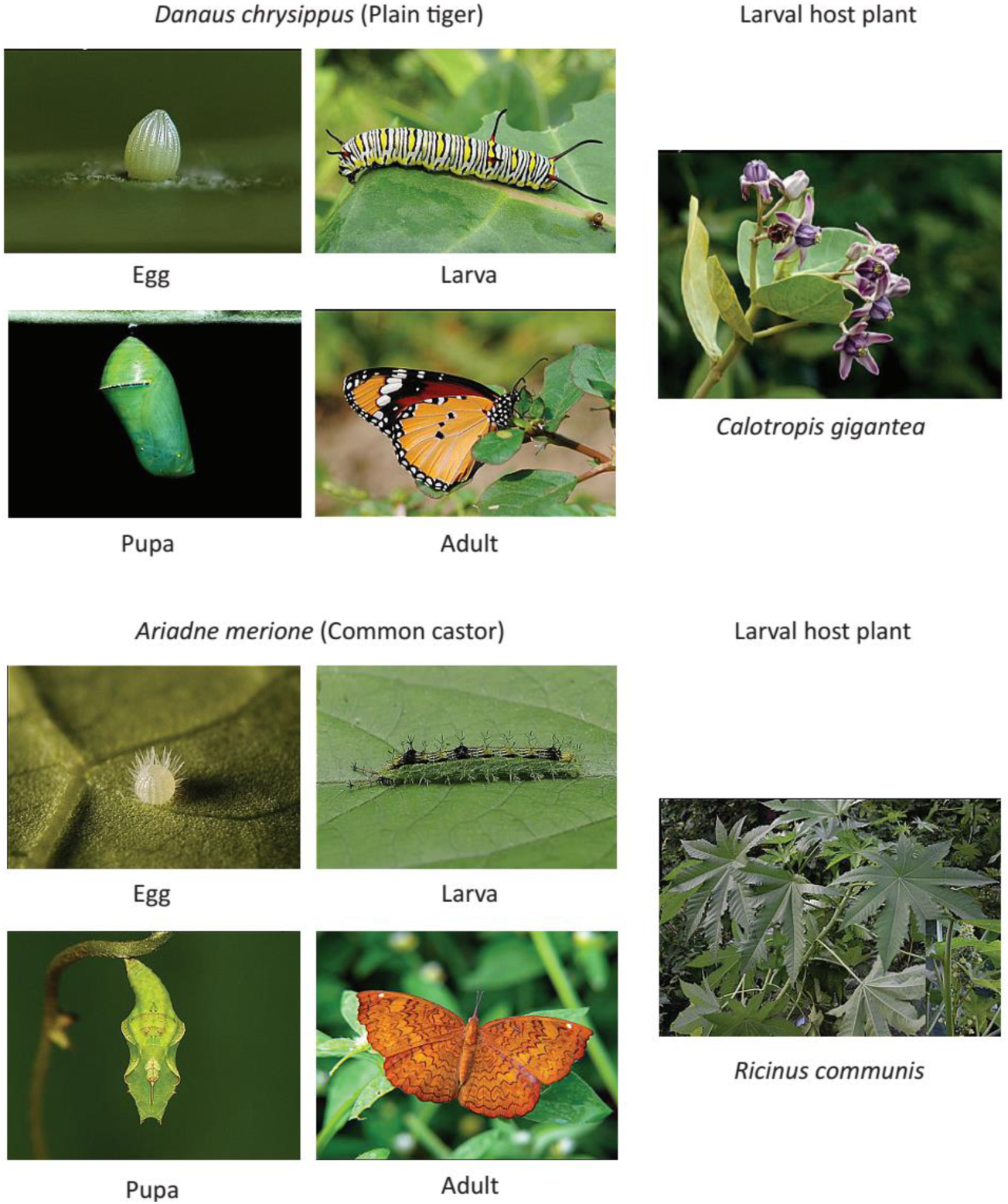
Life stages and larval host plants of *Danaus chrysippus* and *Ariadne merione.* Pictures of key developmental stages of focal butterfly species and larval host plants that were used in this study. Images are sourced from http://www.ifoundbutterflies.org

We eliminated gut microbes from *D. chrysippus* and *A. merione* larvae using two independent methods: a) chemically sterilizing the larval diet, and b) administering antibiotics through the larval diet. Importantly, we conducted these experiments with wild-caught insects and naturally available larval host plants to maximize the relevance of our results to natural insect populations. We tested whether eliminating gut microbes affected the development and survival of *D. chrysippus* and *A. merione.* We found that disrupting gut bacteria did not affect butterfly growth and survival, suggesting the general absence of strong associations between butterflies and their gut bacterial communities.

## MATERIALS AND METHODS

### Insect collection and rearing

To test the impact of removing or disrupting the bacterial community from larval guts, we conducted experiments in multiple blocks for each host species. In each block, we split butterfly eggs into the following broad treatments: (a) control, with no external treatment of leaves, (b) larvae fed with antibiotics or chemically sterilized leaves, and (c) re-introduction of microbes from the larval gut or from leaf surfaces on previously sterilized leaves. We collected *D. chrysippus* males and females on the campus of the National Centre for Biological Science (NCBS) (13.0716° N, 77.5794° E). We kept adult butterflies in cages (height ∼2ft, length ∼1ft and width ∼1ft) along with their host plant *C. gigantea* in a climate-controlled greenhouse maintained at 27-31°C. In each cage, we kept 1 female and ∼1-2 males (in case the female had not mated in the wild), along with nectaries containing artificial nectar solution (Birds Choice #NP1005, USA). From each female, we obtained ∼20–80 eggs, which we distributed equally into different treatment groups. We obtained *D. chrysippus* eggs in the greenhouse because we found very few eggs on larval host plants in nature. On the other hand, we found relatively large numbers of *A. merione* eggs on *R. communis* plants around NCBS, which we used directly. For each block, we collected *A. merione* eggs from 5–10 host plants. Note that the number of replicate larvae in each block and treatment was variable, and depended on the number of eggs that were available. For each host species, we placed individual eggs in small plastic containers (height ∼ 5cm and diameter ∼ 5cm) that were maintained in a larger plastic box (height ∼12cm, length ∼ 1ft and width ∼20cm). Every 24–48 hours, we supplied larvae with fresh leaves collected from the natural habitat. We used leaves from 3–7 different host plants to include variation across plants and associated microbial communities.

### Chemical sterilization of diet

We carried out all experimental procedures in a laminar hood to minimize contamination by environmental microbes. To eliminate microbes from *C. gigantea* leaves, we dipped them in 70% ethanol for 60 seconds and 10% bleach for 30 seconds, followed by three washes with sterile distilled water. We dried leaves completely and cut them into smaller pieces before feeding the larvae. To disentangle the effects of sterilizing agents and microbial elimination, we re-introduced larval gut flora and leaf flora on pre-sterilized leaves using two additional treatments. In one treatment, we created a frass (larval excreta) solution by suspending ∼500 mg frass from control group larvae in 5 ml of sterile Phosphate-buffered saline (PBS). Control group larvae fed on untreated leaves that were expected to harbor the natural microbial community. In the second treatment, we swabbed leaf surfaces of wild *C. gigantea* leaves and suspended the swabs in 5 ml sterile PBS. We painted frass or leaf swab solutions on one side of chemically sterilized leaves, and allowed the leaf surface to dry before feeding larvae. We did not chemically sterilize *R. communis* leaves because they became limp and permanently lost form when dipped in ethanol and bleach. Hence, we only used antibiotic treatment for *A. merione* larvae, as described below.

### Antibiotic treatment

We administered two doses of antibiotics to *D. chrysippus* and *A. merione* larvae. The low dose treatment consisted of a mixture of Ampicillin (500 µg/ml), Tetracycline (50 µg/ml) and Streptomycin (100 µg/ml) in sterile water, and the high dose treatment contained twice as much of each antibiotic. We selected antibiotic concentrations based on previous studies with other insects that reported a significant reduction in gut bacteria [6,8,32,33]. We applied the antibiotic cocktail on both sides of leaves. For *D. chrysippus*, in two out of four experimental blocks, we painted the antibiotic solution on the leaves using a sterile paintbrush; for the other two blocks, we sprayed the antibiotic solution on leaves. For *A. merione*, we sprayed the antibiotic solution on leaves in all blocks. Each spray delivered 150-200 µl antibiotic solution; we sprayed each side of each leaf 4-6 times. As a solvent control, we painted or sprayed leaves with sterile double-distilled water. We let leaf surfaces dry before feeding larvae, administering antibiotics with every feeding (every 24-48 hrs.) until pupation.

### Determining larval gut flora

To quantify the degree of disturbance in bacterial communities of larvae fed with antibiotics and sterile diet, we sequenced the bacterial 16S rRNA gene on an Illumina MiSeq platform, at our in-house sequencing facility. We extracted DNA from larvae from control and treated groups (n=2-3) using a Wizard genomic DNA extraction kit (Promega) and amplified the V3-V4 hypervariable region of the 16S rRNA gene using 300 bp paired-end sequencing as per the standard Illumina MiSeq protocol [34]. We tested for, but did not find evidence of, contamination from DNA extraction kits (see supplementary methods). We analyzed de-multiplexed sequences using QIIME (version 1.9.1; see supplementary methods) [35]. We filtered reads for quality using a minimum quality score of q30 and removed chimeric sequences using USEARCH (version 6.1) [36]. We assembled filtered reads into Operational Taxonomic Units (OTUs) with 97% sequence similarity using UCLUST, with the ‘open-reference OTU picking’ method in QIIME. To determine taxonomy, we compared one representative sequence from each OTU against the Green Genes 16S ribosomal gene database (Greengenes Database Consortium, version gg_13_5) using default QIIME parameters. We used Permutational multivariate ANOVA (permanova, Adonis, package “Vegan”) [37] in R [38] to compare bacterial communities of treated and untreated larvae.

To visualize the differences in bacterial communities across larvae with intact (control) vs. perturbed (treated) gut flora, we carried out ordination analysis of bacterial communities based on bacterial abundance and composition. We tested whether control and treated samples clustered differently using both constrained and unconstrained ordination analysis. Unconstrained ordination analyzes samples without any *a priori* information about groups (e.g. control vs. treated), whereas in constrained ordination, sample groups are pre-defined. We performed Principle Component Analysis as unconstrained ordination using the package “pca3d” in R [39]. For constrained ordination we performed Canonical Analysis of Principal Coordinates based on discriminant analysis (CAPdiscrim) using the R package “BiodiversityR” [40].

To validate MiSeq results, we performed quantitative PCR (qPCR) to compare the amplification of bacterial 16S rRNA genes from dominant bacterial groups across control and treated samples. For qPCR, we used the same DNA samples that we used for MiSeq analysis, quantifying the abundance of Gammaproteobacteria and Actinobacteria relative to an internal control (18S rRNA gene of the host butterfly; see supplementary methods).

### Measuring host fitness and statistical analysis

For each host species, we conducted experiments in 3-4 blocks and measured 4-7 fitness proxies in each case (see supplementary tables S1-S3 and supplementary methods), from the time eggs hatched until adults eclosed. We measured larval length (throughout development), larval weight, pupal weight, time taken from hatching until pupation, time taken from pupation until eclosion, and the weight of freshly eclosed adults. For some experimental blocks, we also estimated larval digestion efficiency by measuring the gain in larval weight per unit time and per gram of leaf consumed, and the amount of excreta produced by larvae per gram of leaf consumed. We tested whether each fitness parameter differed significantly across different treatments using generalized linear models (GLM), followed by Tukey’s post hoc test for multiple comparisons in R, package “multcomp” [41]. For comparisons that were significant after performing Tukey’s multiple comparison test, we report p values and log odds – “estimate (E)”. Finally, we tested whether larval survival varies across treatments using Fisher’s exact test in R. We carried out pairwise comparisons of larval mortality across control and treated groups (for instance, untreated leaves vs. sterilized leaves) in each block to test whether bacterial elimination affected larval survival.

Overall, we tested whether removing gut bacteria altered larval growth, resource use, time required for metamorphosis and larval survival. We could not test the impact of removing bacteria on adult fitness because logistically it was very challenging to maintain large numbers of adults in sterile conditions during mating and oviposition. Furthermore, since prior work shows that pupal and adult weights are good predictors of fecundity in lepidopterans [42–46], we could obtain indirect estimates of adult fitness from these measures.

## RESULTS

### Antibiotic treatment and dietary sterilization effectively disrupted larval microbiomes

Both dietary sterilization and antibiotic treatment significantly altered bacterial communities across treated and untreated groups (Permutational multivariate ANOVA, p<0.05, 10000 permutations, figure 2, panels A1-C1 and A2-C2; also see figures S1 & S2). Treated larvae also had lower bacterial loads, reflected in the increased relative abundance of chloroplast and mitochondrial reads compared to untreated larvae (figure 2, panels A3-C3; a reduction in the number of bacterial 16S gene copies allows higher amplification of leaf-derived chloroplast and leaf or host-derived mitochondria). Finally, quantitative PCR confirmed that bacterial load reduced dramatically after antibiotic administration and dietary sterilization (figure S3).

**Figure 2.**
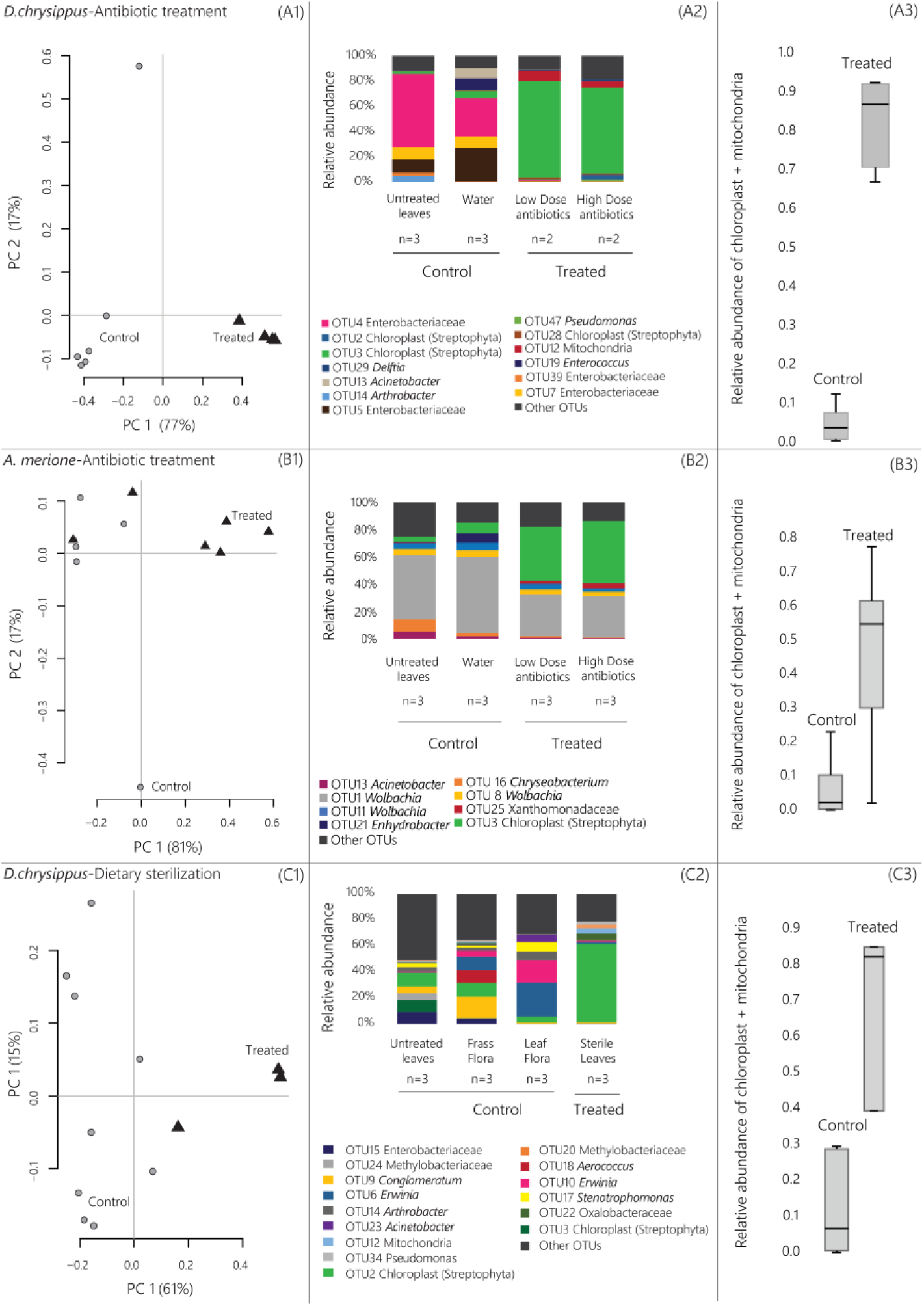
Effect of antibiotic treatment and dietary sterilization on bacterial communities of *D. chrysippus* and *A. merione* larvae. Panels (A1-C1): Stacked bar plots show the mean relative abundance of the five most abundant bacterial taxa (OTUs, identified to the lowest taxonomic level possible) in each treatment group (see supplementary methods). Panels (A2-C2): Boxplots (median with quartiles; whiskers show data range) show the relative abundance of reads assigned to chloroplasts and mitochondria (combined) in control and treated larvae. Panels (A3-C3): Principle component analysis (PCA) of full bacterial communities from control and treated larvae. Axes show the first two principle components (PC) that explain maximum variation in the data; values in parenthesis show the percent variation explained by each PC. For microbiome analysis, we used *D. chrysippus* larvae from block 2 (dietary sterilization) and block 3 (antibiotic treatment), and *A. merione* larvae from block 1 (antibiotic treatment) (see supplementary tables S1-S3 and figures 4-6).

### Dietary sterilization does not impact the fitness of *D. chrysippus*

We started our manipulative experiments with *D. chrysippus,* feeding larvae with surface sterilized *C. gigantea* leaves. In our first experimental block (block 1) we found that larvae fed on sterile diet grew more slowly than the control group and pupated ∼48 hours later (figure 3A-C; GLM, model: fitness ∼ treatment; Tukey’s post hoc test for multiple comparisons, p < 0.05; table S1). However, we did not observe any larval mortality (table 1); and pupal weight, adult weight and time taken for eclosion (pupal span) remained unaffected (figure 3D-F). To confirm that slow larval growth occurred due to bacterial elimination and not due to toxicity from sterilizing chemicals, in two subsequent blocks (block 2 and block 3) we re-introduced natural microflora on sterilized *C. gigantea* leaves (see methods), for a total of four treatment groups – 1) non-sterile (untreated) leaves 2) sterile leaves, 3) sterile leaves with frass flora and 4) sterile leaves with leaf flora. In block 2 and block 3, for most fitness measurements, we did not find significant differences across treatments (figure 4 and figure S4; GLM model: fitness ∼ treatment, Tukey’s post hoc test for multiple comparisons, p > 0.05; table S1). However in a few cases, fitness of treated individuals (fed with sterile leaves) was lower than control (fed with untreated leaves). For instance, adult weight in block 2 (figure 4, panel E) and larval span in block 3 (figure S4, panel B) were significantly affected in treated individuals (GLM, model: fitness ∼ treatment, Tukey’s post hoc test for multiple comparisons, p <0.05). However, these effects were neither consistent across blocks nor across fitness measurements (table S1). Moreover, in these cases, there was no significant variation in fitness across treated individuals (fed with sterile leaves) and individuals with re-introduced microbiota (figure 4 and S4, table S1). Adding bacteria from frass or leaf surfaces also did not affect larval growth, except in one case (larval span, block 3, figure S4, panel B). Similarly, larval mortality was not significantly different across treatment groups (Fisher’s exact test, p>0.05, table 1). Together, these results show that feeding surface-sterilized diet to *D. chrysippus* larvae had weak and variable impacts on host fitness.

**Table 1:** Larval mortality across experimental blocks and treatments. We tested for pairwise differences between survival of control (untreated leaves or leaves + water) and treatment groups separately for each block using a Fisher’s exact test. *DC* and *AM* represent *D. chrysippus* and *A. merione* respectively. The only significant difference was found in *A. merione* block 3 (p=0.02*), where mortality in (Leaves + Water) < (Leaves +High Dose antibiotic).

| Dietary sterilization | Untreated leaves |  | Sterile leaves |  | Sterile leaves + leaf flora |  | Sterile diet + frass flora |  | p Value for Fisher's exact test |
| --- | --- | --- | --- | --- | --- | --- | --- | --- | --- |
|  | Total number of larvae | % Dead larvae | Total number of larvae | % Dead larvae | Total number of larvae | % Dead larvae | Total number of larvae | % Dead larvae |  |
| *DC* block 1 | 9 | 0% | 9 | 0% | -- | -- | -- | -- | No Mortality |
| *DC* block 2 | 13 | 0% | 15 | 7% | 12 | 0% | 13 | 23% | $p>0.05$ |
| *DC* block 3 | 11 | 18% | 13 | 23% | 11 | 0% | 11 | 18% | $p>0.05$ |
| Antibiotic treatment | Untreated leaves |  | Leaves + Water |  | Leaves + Low dose antibiotics |  | Leaves + High dose antibiotics |  | p Value for Fisher's exact test |
| *DC* block 1 | 5 | 0% | 5 | 0% | 8 | 0% | 8 | 0% | No Mortality |
| *DC* block 2 | 14 | 7% | 16 | 6% | 15 | 7% | 18 | 0% | $p>0.05$ |
| *DC* block 3 | 29 | 10% | 26 | 4% | 23 | 4% | 25 | 4% | $p>0.05$ |
| *DC* block 4 | 13 | 0% | 13 | 0% | 11 | 27% | 13 | 0% | $p>0.05$ |
| *AM* block 1 | 18 | 6% | 17 | 24% | 19 | 26% | 19 | 21% | $p>0.05$ |
| *AM* block 2 | 11 | 36% | 8 | 25% | 12 | 0% | 11 | 9% | $p>0.05$ |
| *AM* block 3 | 10 | 20% | 8 | 13% | 9 | 33% | 10 | 70% | $p<0.05^*$ |

**Figure 3.**
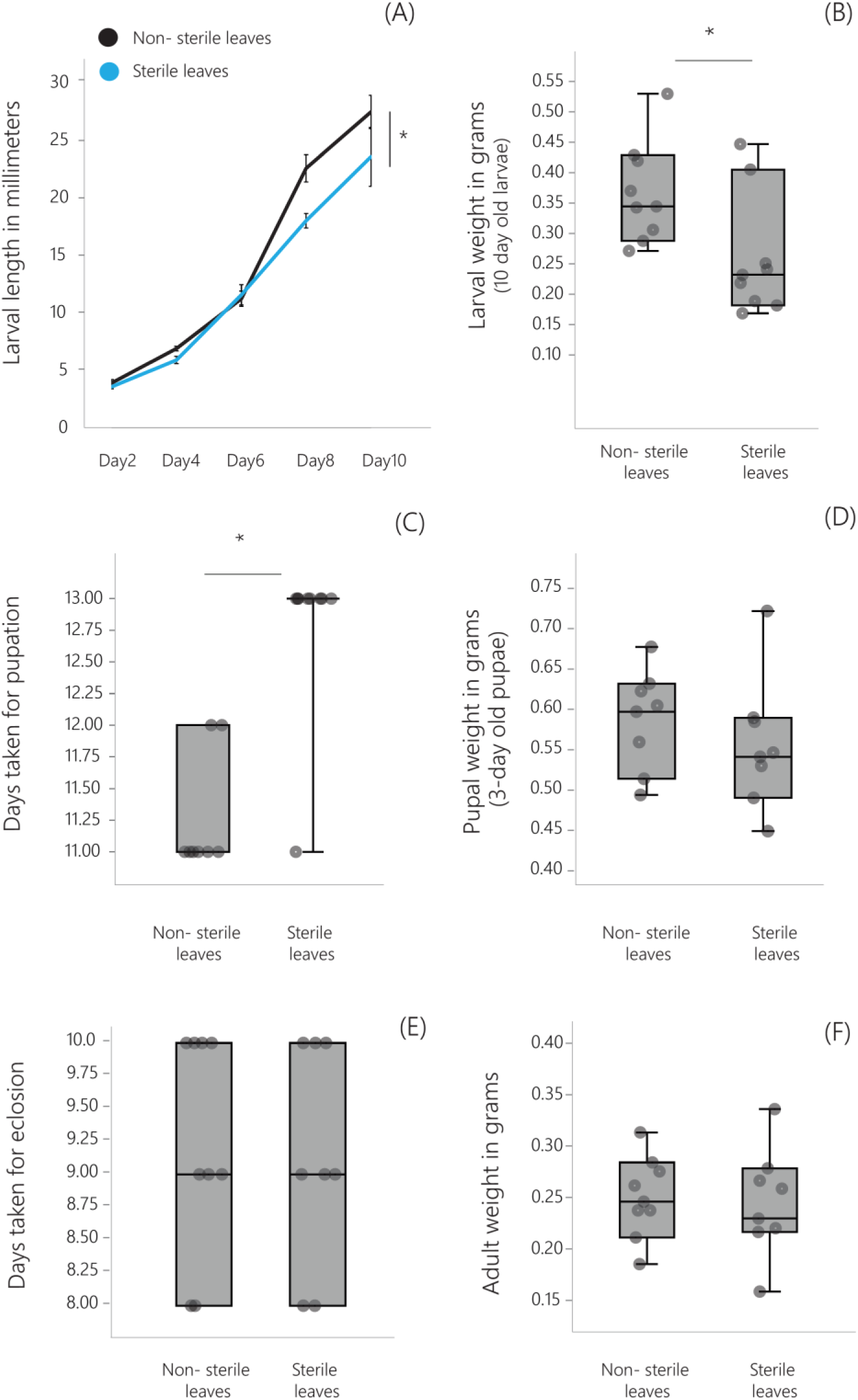
Effect of dietary sterilization on *D. chrysippus* fitness. Panels show different fitness measurements for experimental block 1 (results from other blocks are shown in figures 4 and S4). Asterisks indicate a significant difference between control and treatment groups (GLM, model: fitness ∼ treatment, Tukey’s post hoc test for multiple comparisons, p < 0.05). For each treatment group, n=∼8 individuals (see table S1).

**Figure 4.**
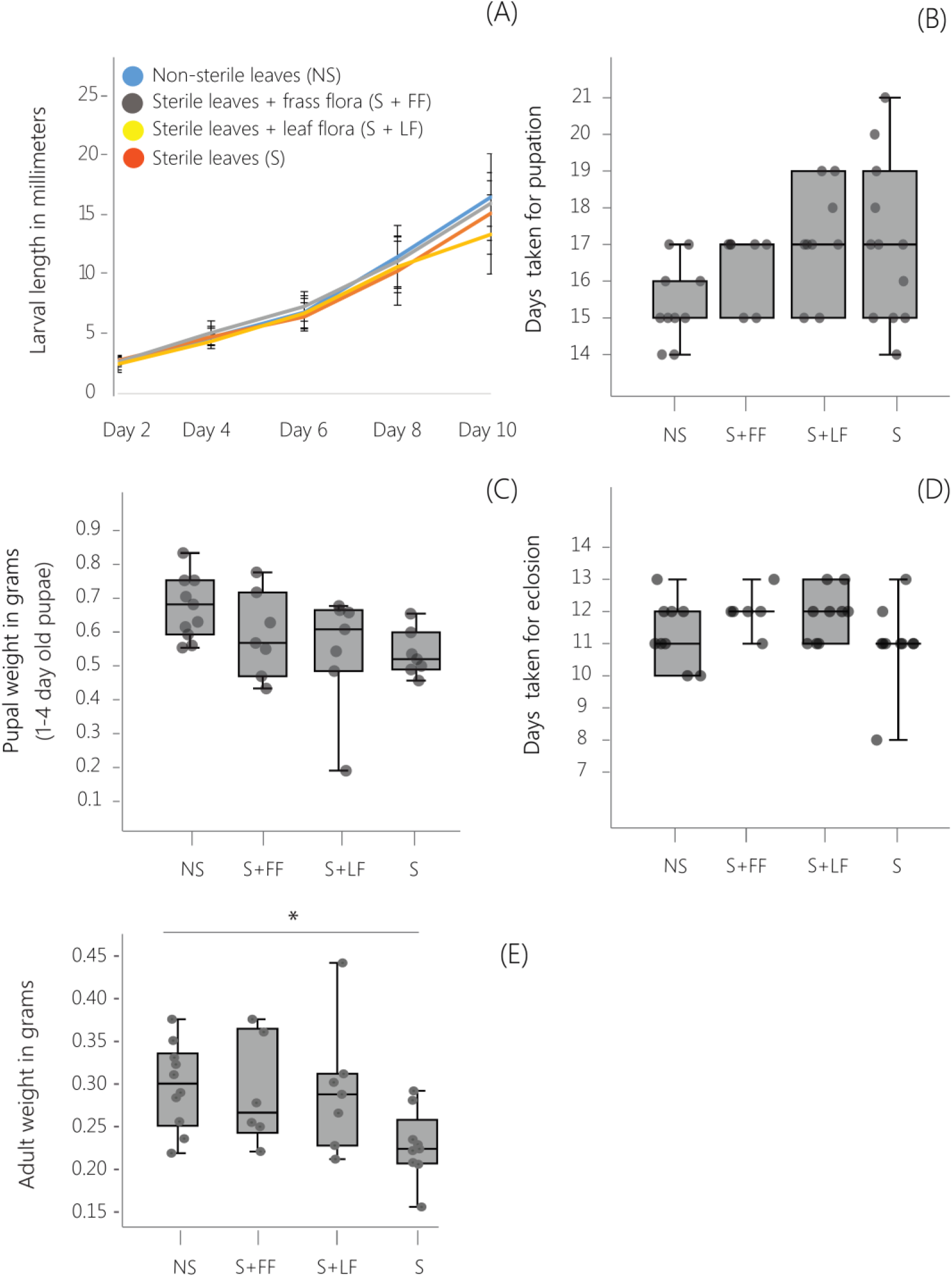
Effect of dietary sterilization and microbial re-introduction on *D. chrysippus* fitness. Panels show different fitness measures for experimental block 2 (results from other blocks are shown in figure S4 and table S1). Asterisks indicate a significant difference between treatment groups (GLM, Tukey’s test for multiple comparisons, p < 0.05). For each treatment group, n= 6-13 individuals (see table S1).

### Antibiotic treatment does not impact the fitness of *D. chrysippus* and *A. merione*

Though surface sterilization of diet effectively eliminates leaf surface microbes, it may not eliminate bacteria that reside within plant tissues or on the egg casing (which is sometimes consumed by larvae; [47]). To eliminate bacteria from these sources, we fed larvae of *D. chrysippus* and *A. merione* with a cocktail of broad-spectrum antibiotics, and tested the impact on their growth. For both host species, we did not observe a significant difference in fitness proxies across control (leaves + water) and treated (leaves + antibiotics) individuals (figures 5 & 6; figures S5-S8; GLM, model: fitness ∼ treatment, Tukey’s post hoc test for multiple comparisons, p > 0.05; tables S2 & S3), except in a few cases (see figures 5,6,S5-S8 and tables S2 & S3). Again, the observed effect was neither consistent across different fitness measurements nor across experimental blocks (see figures S5-S8 and tables S2 & S3). Fitness did not vary across individuals fed with untreated leaves and leaves sprayed with water, except for two fitness measures (pupal and adult weight) in block 2 for *A. merione* (figure S8, panels D and F, see table S3). However, for one of these measurements (adult weight), fitness was not significantly different across leaves sprayed with water and leaves sprayed with antibiotics (see table S3), again suggesting that bacterial elimination does not affect host fitness strongly. Finally, we did not find a significant difference in larval mortality across control and antibiotic treated groups (Fisher’s exact test, p>0.05, Table 1), except in *A. merione* block 3 (p<0.05). However, mortality was only observed in larvae that were fed with high dose of antibiotics but not with a low dose (table1), suggesting that larval death was not consistently associated with antibiotic treatment (and its impact on the larval microbiome).

**Figure 5.**
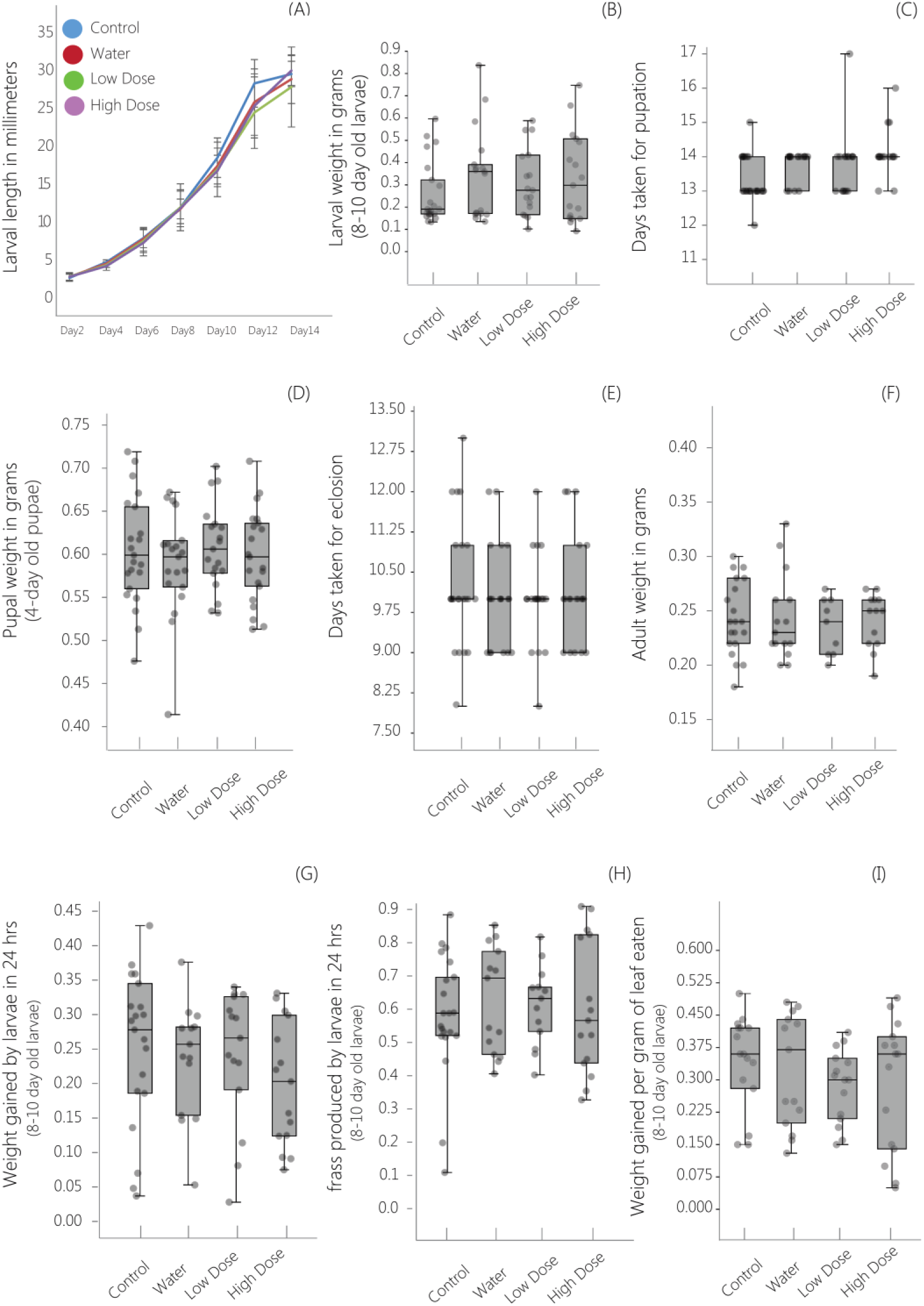
Effect of antibiotic administration on *D. chrysippus* fitness. Panels show different fitness measures for experimental block 3 (results from other blocks are shown in figures S5-S7 and table S2). We did not observe a significant treatment effect for any measurement (GLM, model: fitness ∼ treatment, Tukey’s post hoc test for multiple comparisons, p>0.05). For each treatment group, n= 9-25 individuals (see table S2).

**Figure 6.**
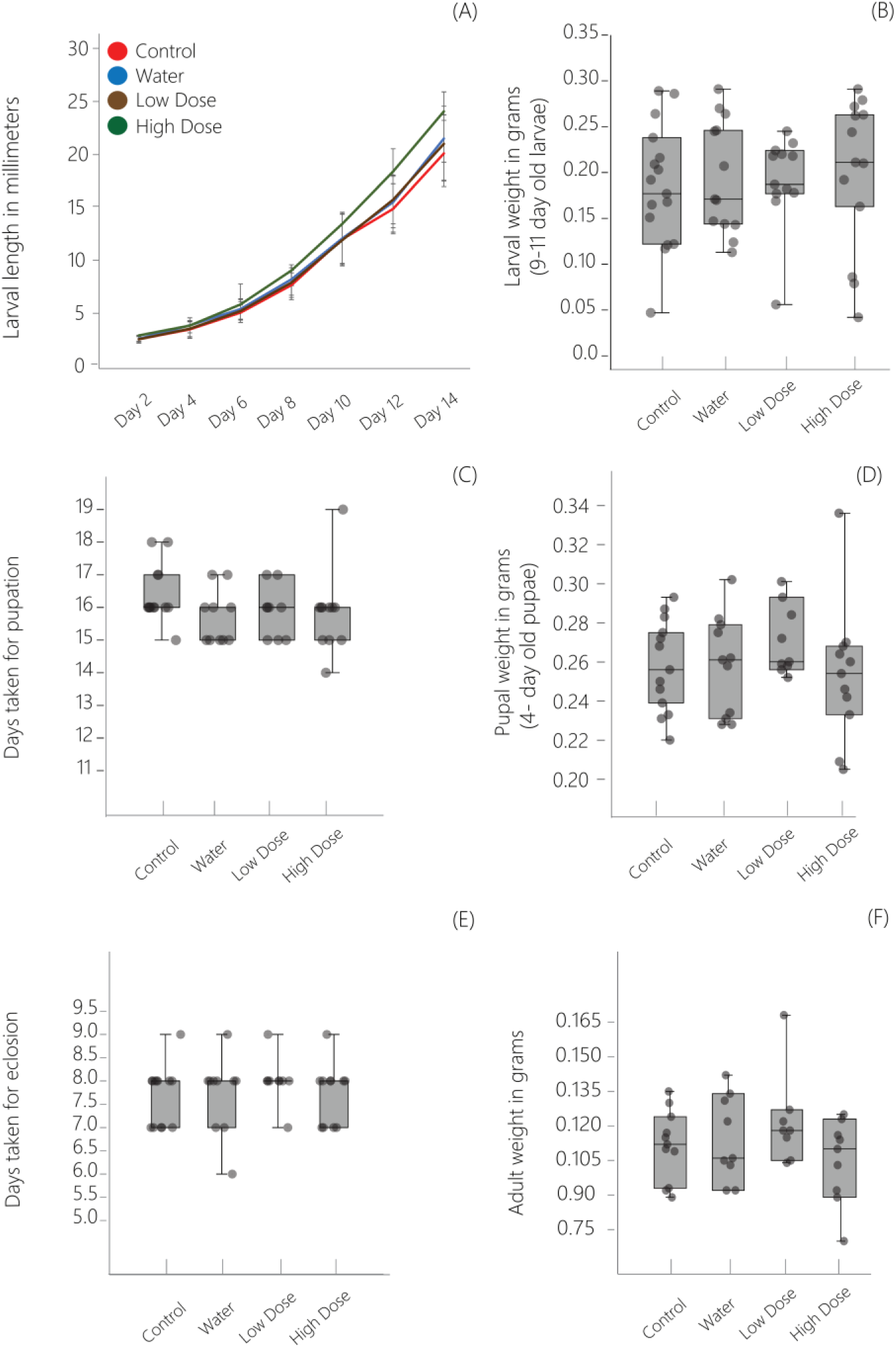
Effect of antibiotic administration on the fitness of *A. merione.* Panels show different fitness measures for experimental block 1 (results from other blocks are shown in figure S8 and table S3). We did not observe a significant treatment effect for any measurement (GLM, model: fitness ∼ treatment, Tukey’s post hoc test for multiple comparisons, p>0.05). For each treatment group, n= 9-16 individuals (see table S3).

Overall, we observed that neither dietary sterilization nor antibiotic treatment affected the development and survival of butterfly larvae (summarized in tables 1 and 2). Notably, we observed that adult weight at eclosion – an important predictor of reproductive fitness – was not affected by the reduction in bacterial loads and disruption of gut bacterial communities. Together, these results suggest that gut bacterial communities have generally weak and negligible impacts on the fitness of *D. chrysippus* and *A. merione* larvae.

**Table 2:** Summary of effects of experimental microbial elimination on butterfly hosts. Larval development includes larval length, larval weight and larval span. Dashes indicate blocks where a specific experimental treatment was not included.

| Butterfly host | Experiment | Effect of treatment on fitness proxies |  |  |  |
| --- | --- | --- | --- | --- | --- |
|  |  | Block 1 | Block 2 | Block 3 | Block 4 |
| *D. chrysippus* | Dietary sterilization | Yes<br>(larval development) | — | — | — |
| *D. chrysippus* | Dietary sterilization and microbial re-introduction | No | No | — | — |
| *D. chrysippus* | Antibiotic treatment | No | No | No | No |
| *A. merione* | Antibiotic treatment | No | No | Yes<br>(larval development and survival) | — |

## DISCUSSION

Associations with bacteria impact the fitness of many insects, suggesting that such relationships represent a general phenomenon [1]. Our study contradicts this general pattern, showing that various aspects of butterfly fitness (early growth and survival) are not affected despite a substantial disturbance and reduction in their gut microbial communities. An extensive review on insect-microbe interactions [1] reports numerous cases where microbes significantly impacted insect fitness, with only two cases where microbes had no negative impact on their insect hosts. In this context, our study is one of very few experimental reports showing that some insects do not depend on their gut microbes (also see a recent report on neutrally assembled microbiomes in dragonflies [48]. In conjunction with recent work [13,14] on butterfly-associated bacterial communities (discussed in the Introduction), our experiments strongly support the idea that butterflies have not established key bacterial mutualisms during their evolution. As suggested previously, this lack of host-bacterial mutualism may arise because butterfly gut morphology and physiology may prevent the growth and establishment of microbes [14]. Additionally, butterflies may have evolved a highly efficient and diverse set of digestive enzymes in conjunction with dietary diversification, allowing larvae to digest diverse host plants without relying on their gut microbes [49]. Finally, butterflies might have evolved microbe-independent mechanisms to deal with the challenges of detoxifying poisonous plants. For instance, *D. chrysippus* has evolved resistance to cardiac glycosides found in milkweeds, via mutations in the Na+/K+ pump [50], potentially weakening any selection favoring detoxification by gut bacteria. However, the relative timescales for the evolution of such host-specific, microbe-independent mechanisms is not clear. It is also possible that butterflies have evolved functional associations with microbes in a different, non-dietary context. For instance, butterfly gut bacteria may play an important role in butterfly immune function, as observed in a few other insects [1,51,52]. Alternatively, as demonstrated earlier in diamondback moths, gut bacteria could assist in insecticide resistance [10]. More generally, it is also possible that dependence on the microbiome may have evolved in the context of environmental fluctuations (which would be dampened in greenhouse and laboratory experiments such as ours), or for adult foraging, fecundity and lifespan (which we could not measure here). Nonetheless, our results pose interesting open questions: how did butterflies occupy vastly different dietary niches without recourse to bacterial mutualists, in contrast to the predominant dependence on microbes observed in other insects with similarly diverse diets?

In conclusion, we suggest that the impact of gut microbes on their hosts may range along a continuum from strong to weak dependence or no association. To predict the impact of gut bacteria on insect diversification and evolution, it is important to know how different insects are distributed across this spectrum. Current literature largely represents only one end of the scale, where gut microbes seem to strongly affect their hosts. In this context, our study on wild butterflies presents an interesting contrast to the general trend. It may be interesting to experimentally explore the trends revealed by our work in natural populations of a larger number of butterfly species with contrasting life histories. Hopefully, we can then begin to understand why some insects depend on gut microbes for survival whereas others remain unaffected.

## Authors’ contributions

DA, KK, and KP conceived the project and designed experiments. KP performed the experiments. KK provided greenhouse space and related infrastructural support, and advice on handling butterflies. KP and DA analysed data and wrote the manuscript with input from KK. All authors gave final approval for the manuscript.

## Acknowledgements

We thank members of the Agashe lab for their constructive comments on the manuscript; Arun Prakash, Jagrithi Ramnathan, Jagath Vedamurthy and Sanah Imani for laboratory assistance; members of the Kunte lab for help in the greenhouse and with butterflies; Aparna Agarwal and Rittik Deb for help with 16S MiSeq sequencing and analysis; and Awadhesh Pandit and Tejali Naik from the Next Generation Sequencing (NGS) Facility at NCBS for help with sequencing.

