## Supplementary Information for "Disrupting butterfly microbiomes does not affect host survival and development"

Kruttika Phalnikar, Krushnamegh Kunte and Deepa Agashe
National Centre for Biological Sciences, GKVK Campus, Bellary road, Bangalore, India

### SUPPLEMENTARY FIGURES

**Figure S1- Variation in bacterial communities of larvae across treatments.** Panels show (unconstrained) principle component analysis (PCA) of bacterial communities across experimental treatments. The figure represents the same analysis shown in figure 2; however, here we color each of treatment separately, which are combined in figure 2. Axes represent principle components (PC) explaining maximum variation in the data. Values in parentheses show the proportion of variation explained by each PC. Dietary sterilization and antibiotic administration both significantly affected bacterial communities (Permutational multivariate ANOVA, p<0.05, 10000 permutations).

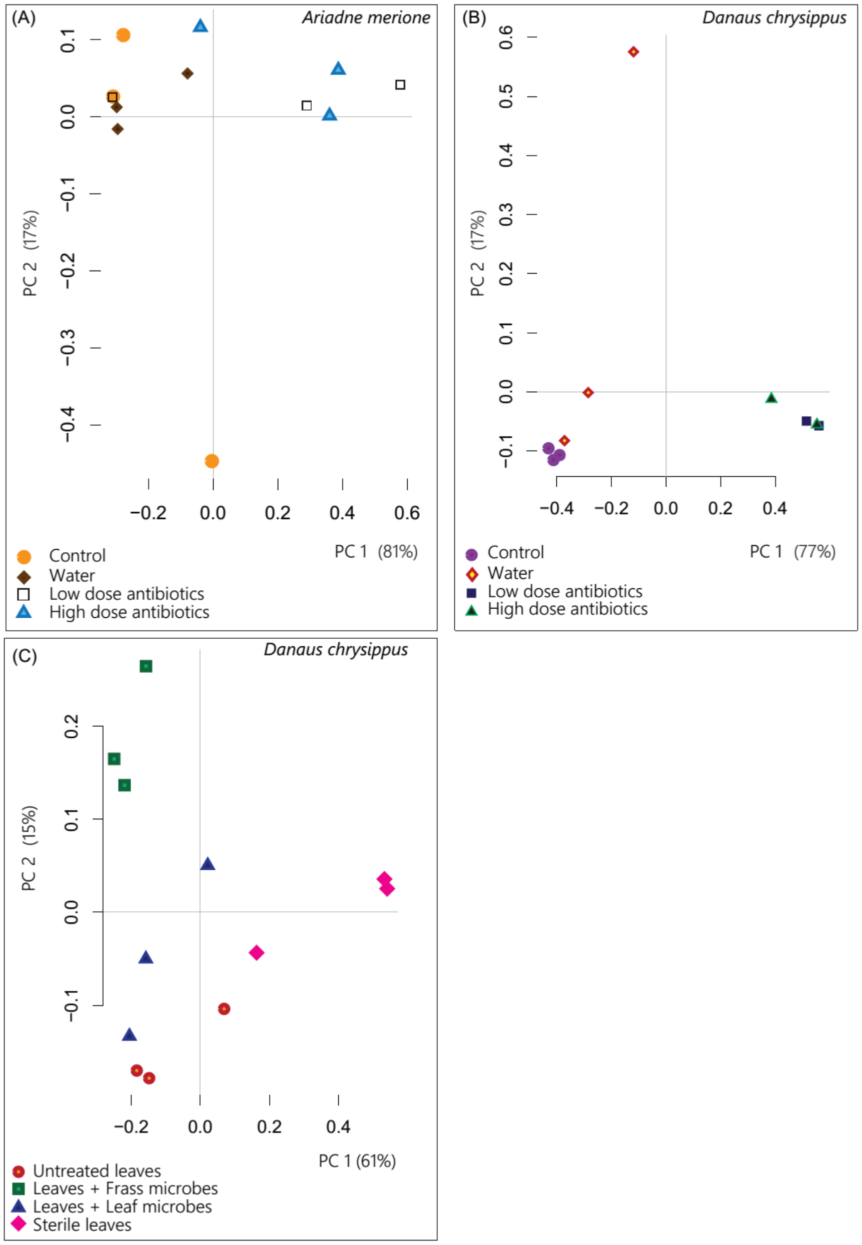

**Figure S2 - Variation in larval bacterial communities across treatments (CAPdiscrim analysis).** Panels show Constrained Analysis of Principal Coordinates (CAP) of communities based on the composition and relative abundance of bacterial OTUs in control and treated groups. Axis labels indicate the proportion of between-group variance (%) explained by the first two linear discriminants (LD1 and LD2). In all panels, LD1 explains 100% between-group variance. Ellipses represent 95% confidence intervals. For each panel, we observed a significant variation in bacterial communities of treated and control group larvae ($p<0.05$, multivariate ANOVA). Larvae with intact and perturbed gut bacterial communities represent control and treated groups, respectively.

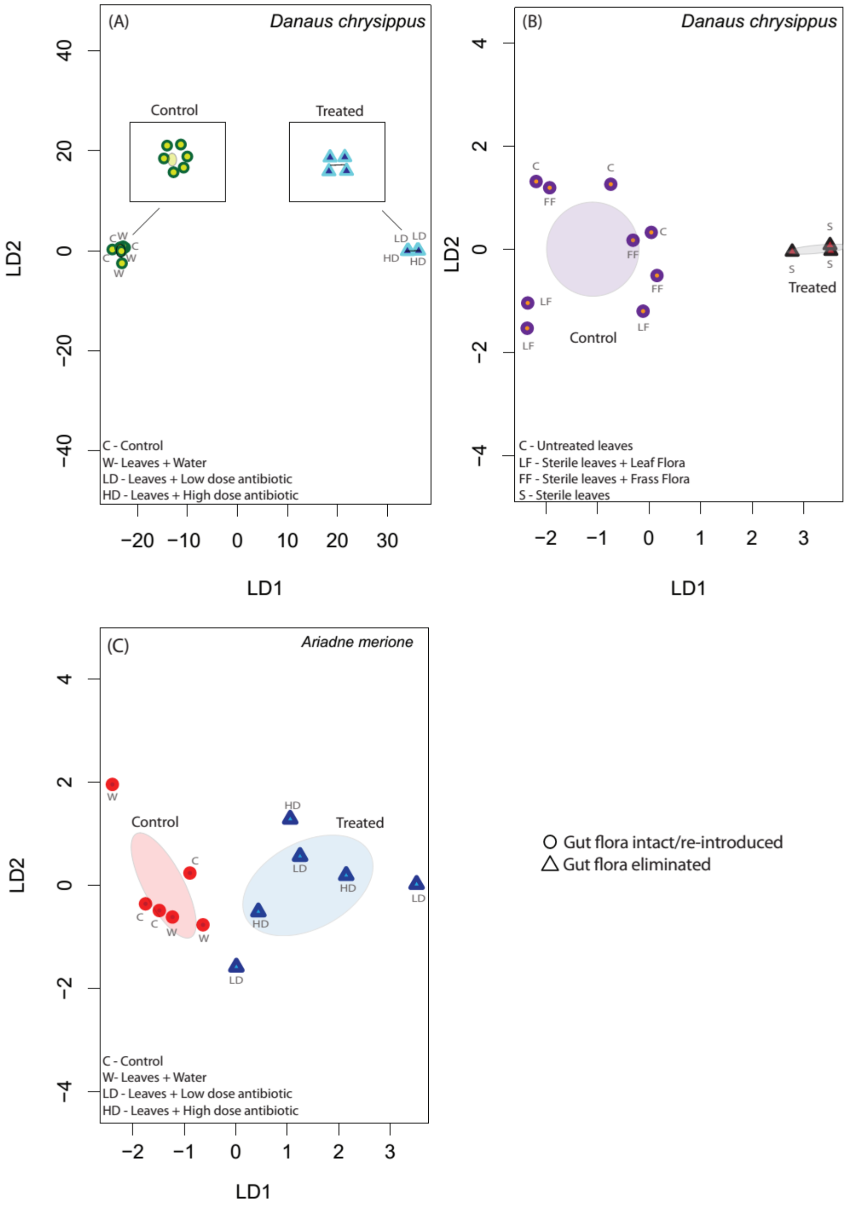

**Figure S3 - Reduction in bacterial abundance after dietary sterilization and antibiotic treatment.** Barplots represent degree of amplification of the target gene (bacterial 16S rRNA gene) relative to an internal control gene (host 18S rRNA gene), calculated as $2^{\wedge(-\Delta CT)}$, where $\Delta CT$ (cycle threshold) = target gene CT – internal control gene CT (n=2-3 larvae). Error bars represent standard deviation. Values in parentheses represent the mean fold reduction in bacterial abundance in treated samples as compared to control samples, calculated as ($2^{\wedge(-\Delta CT)}$ control / $2^{\wedge(-\Delta CT)}$ treated).

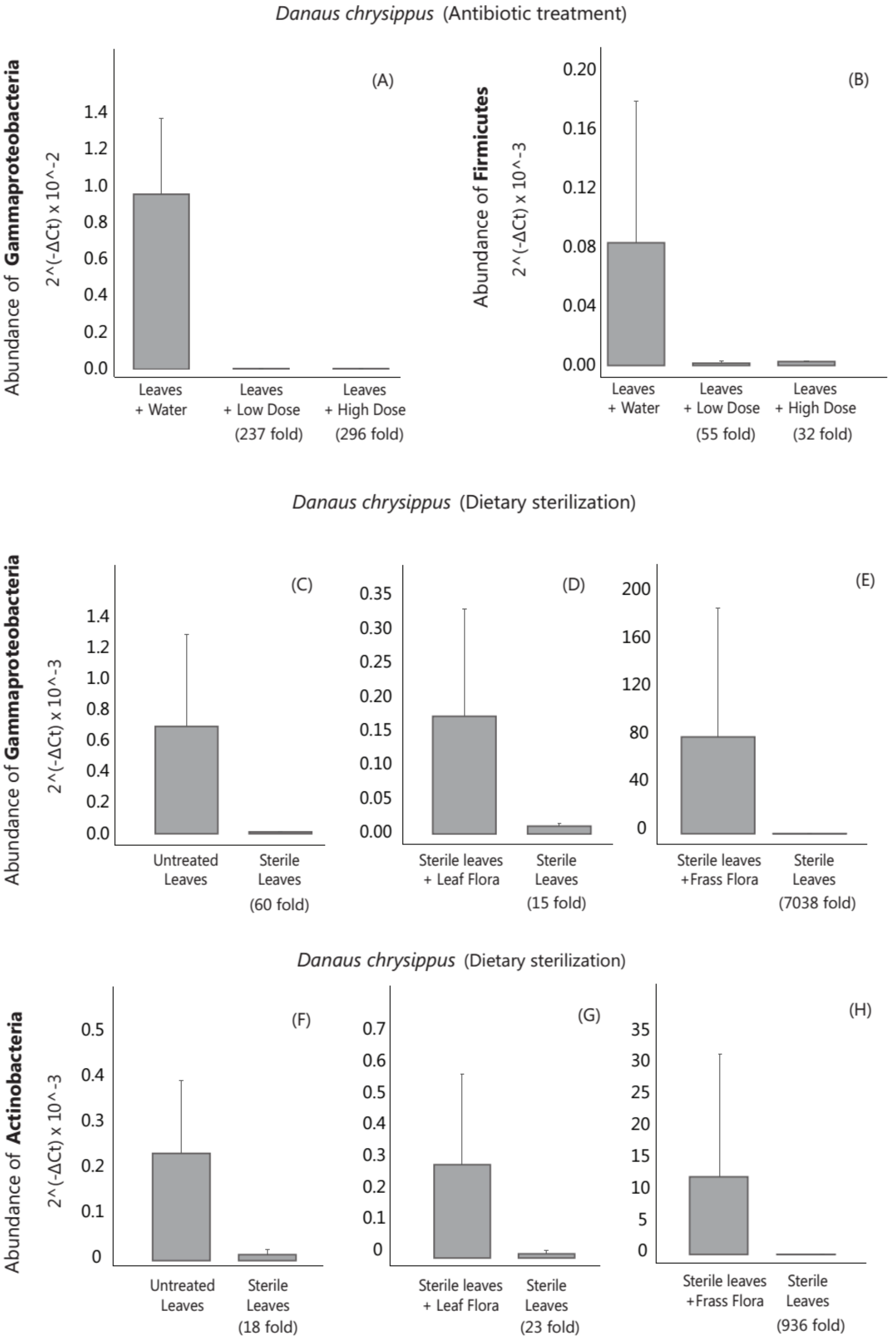

**Figure S4 - Effect of dietary sterilization and microbial re-introduction on *D. chrysippus* fitness.** Panels show different fitness measures for experimental block 3 (results from all 3 blocks for dietary sterilization and microbial re-introduction are summarized in table S1; also see figures 3 and 4). Asterisks indicate a significant difference between treatment groups (GLM, Tukey's post hoc test for multiple comparisons, $p < 0.05$). For each treatment group, n= 6-8 individuals.

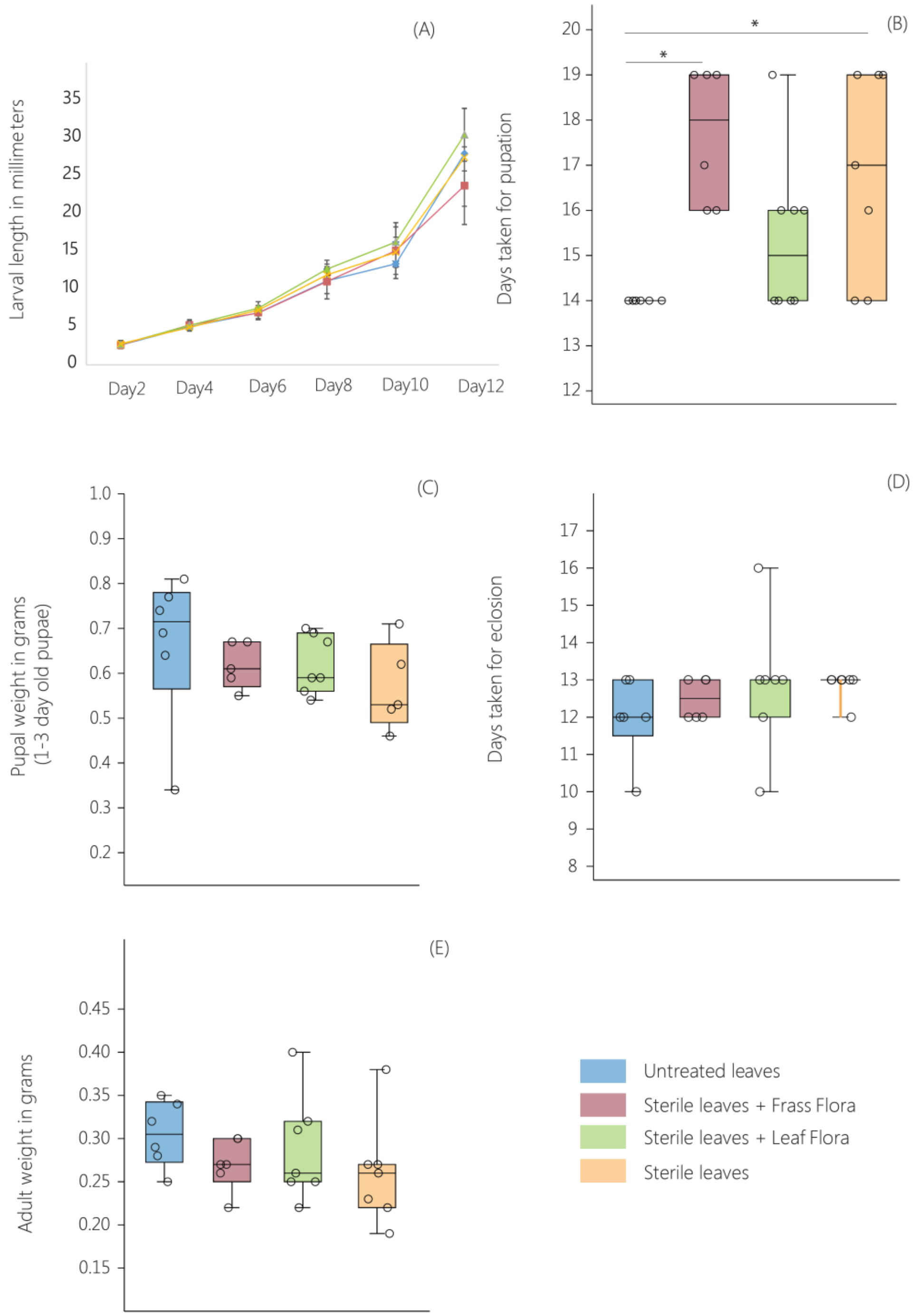

**Figure S5 - Effect of antibiotic administration on *D. chrysippus* fitness.** Panels show different fitness measures for experimental block 1 (results from all 4 experimental blocks are summarized in table S2; also see figure 4 and figures S6-S7). We did not observe a significant treatment effect for any measurement. For each treatment group (GLM, Tukey's post hoc test for multiple comparisons, p >0.05), n= 5-8 individuals.

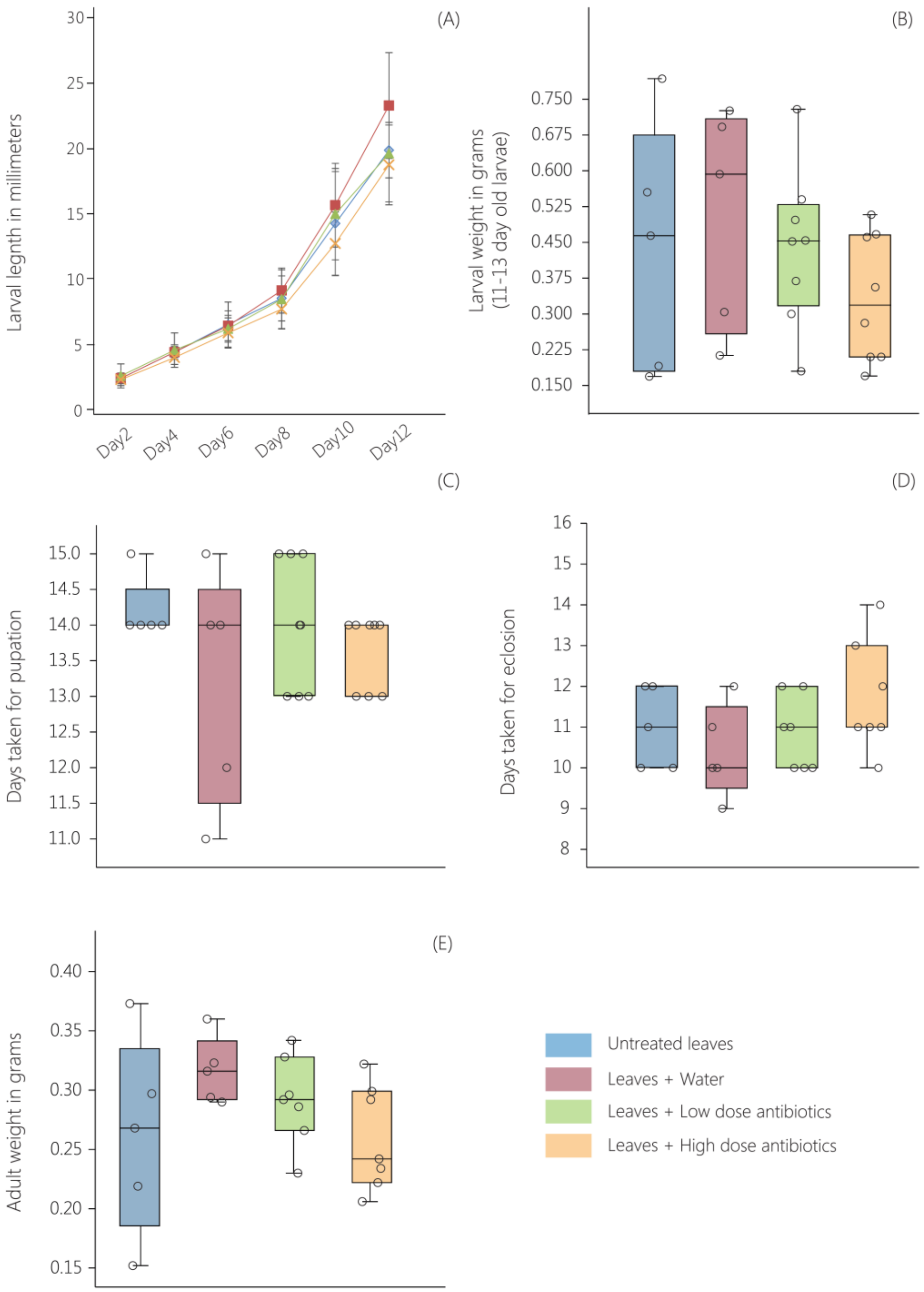

**Figure S6 - Effect of antibiotic administration on *D. chrysippus* fitness.** Panels show different fitness measures for experimental block 2 (results from all 4 experimental blocks are summarized in table S2; also see figures 4, S5 and S7). Asterisks indicate a significant difference between treatment groups (GLM, Tukey's test for multiple comparisons, $p < 0.05$). For each treatment group, n= 4-18 individuals.

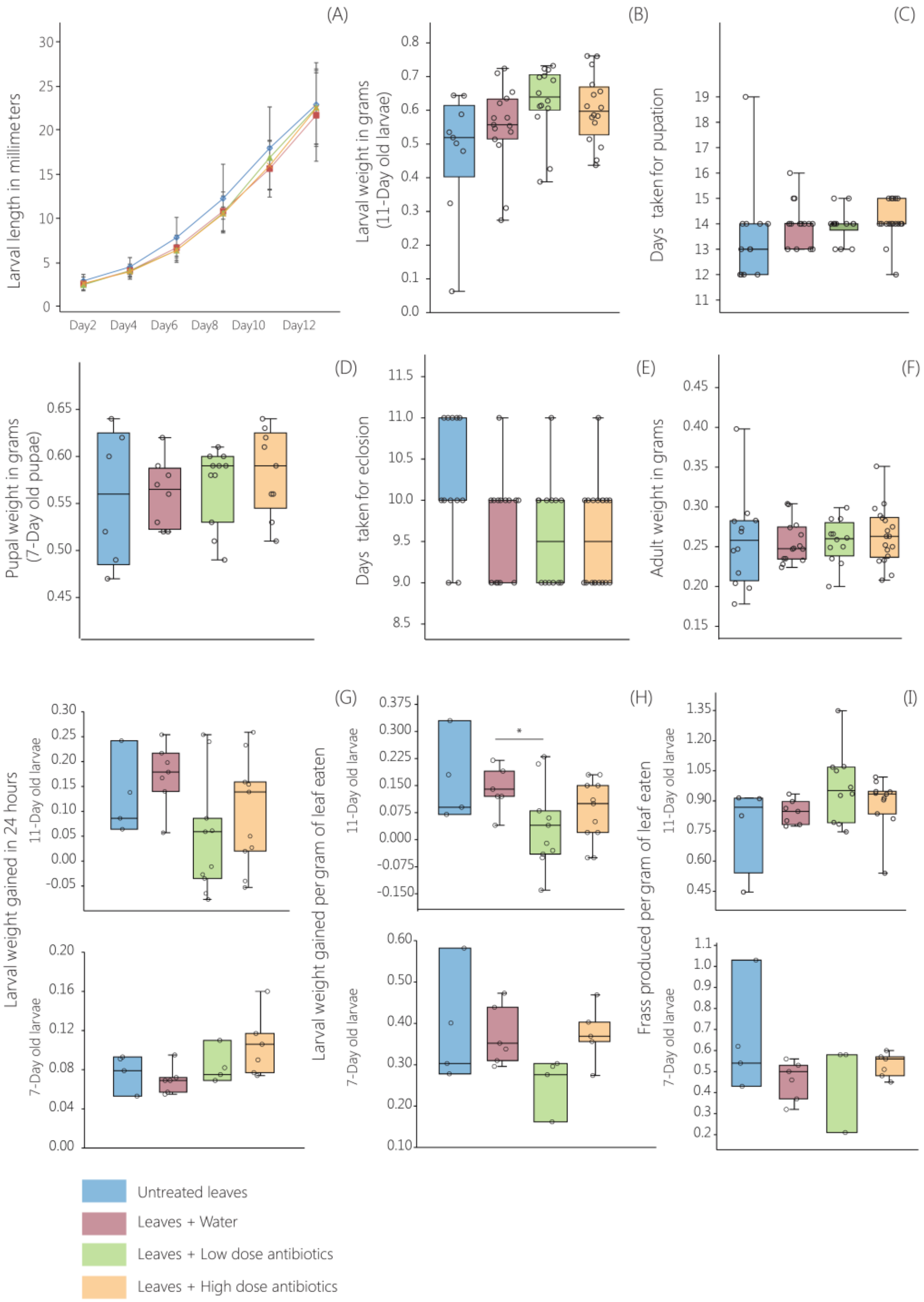

**Figure S7 - Effect of antibiotic administration on *D. chrysippus* fitness.** Panels show different fitness measures for experimental block 4 (results from all 4 experimental blocks are summarized in table S2; also see figures 4, S5 and S6). We did not observe a significant treatment effect for any measurement. For each treatment group, n= 4-13 individuals.

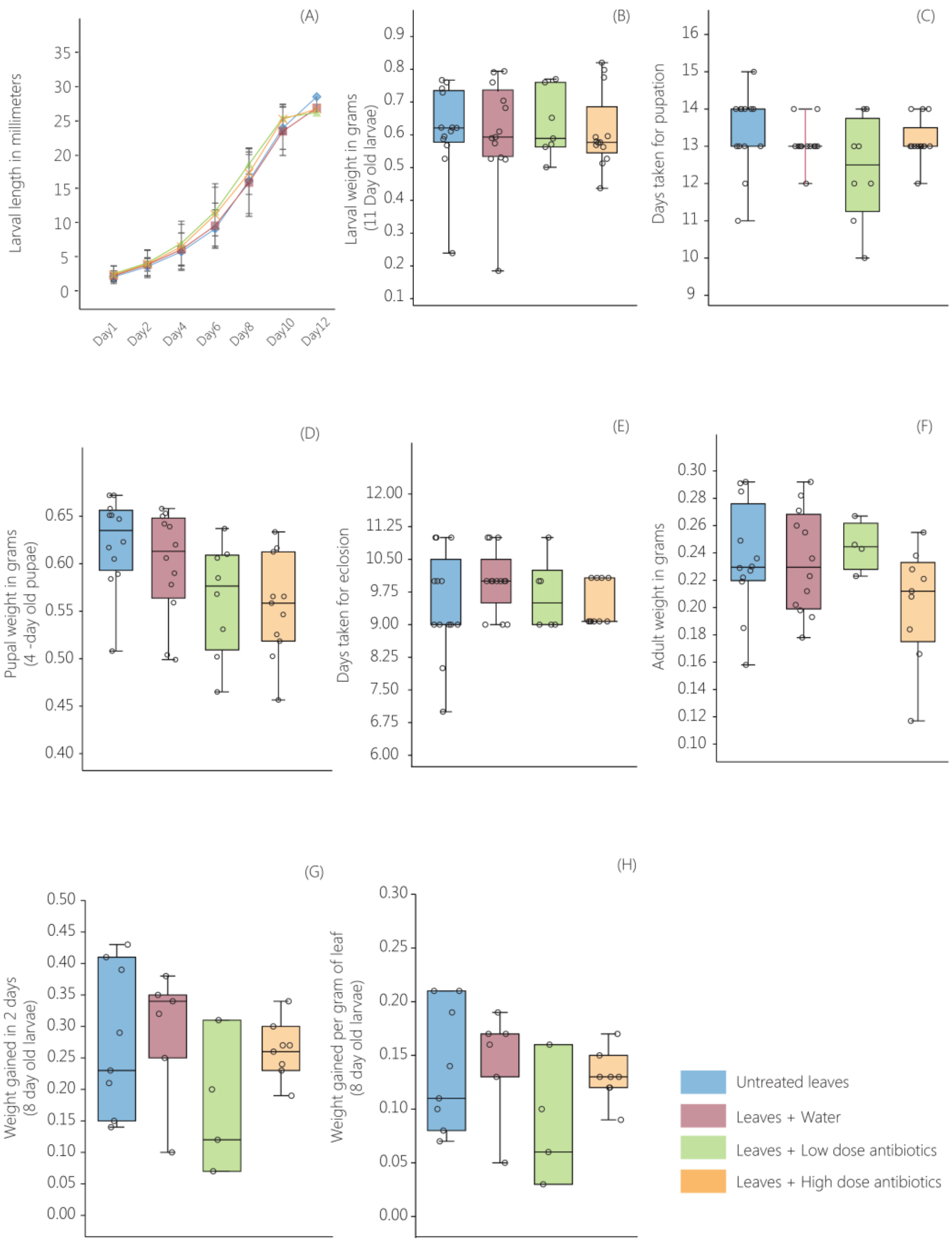

**Figure S8- Effect of antibiotic administration on the fitness of *A. merione*.** Panels show different fitness measures for experimental blocks 2 and 3 (results from all 3 blocks are summarized in table S3). Asterisks indicate a significant difference between treatment groups (GLM, Tukey's test for multiple comparisons, $p < 0.05$). n= 5-12 individuals per treatment for block 2 and n=3-8 for block 3.

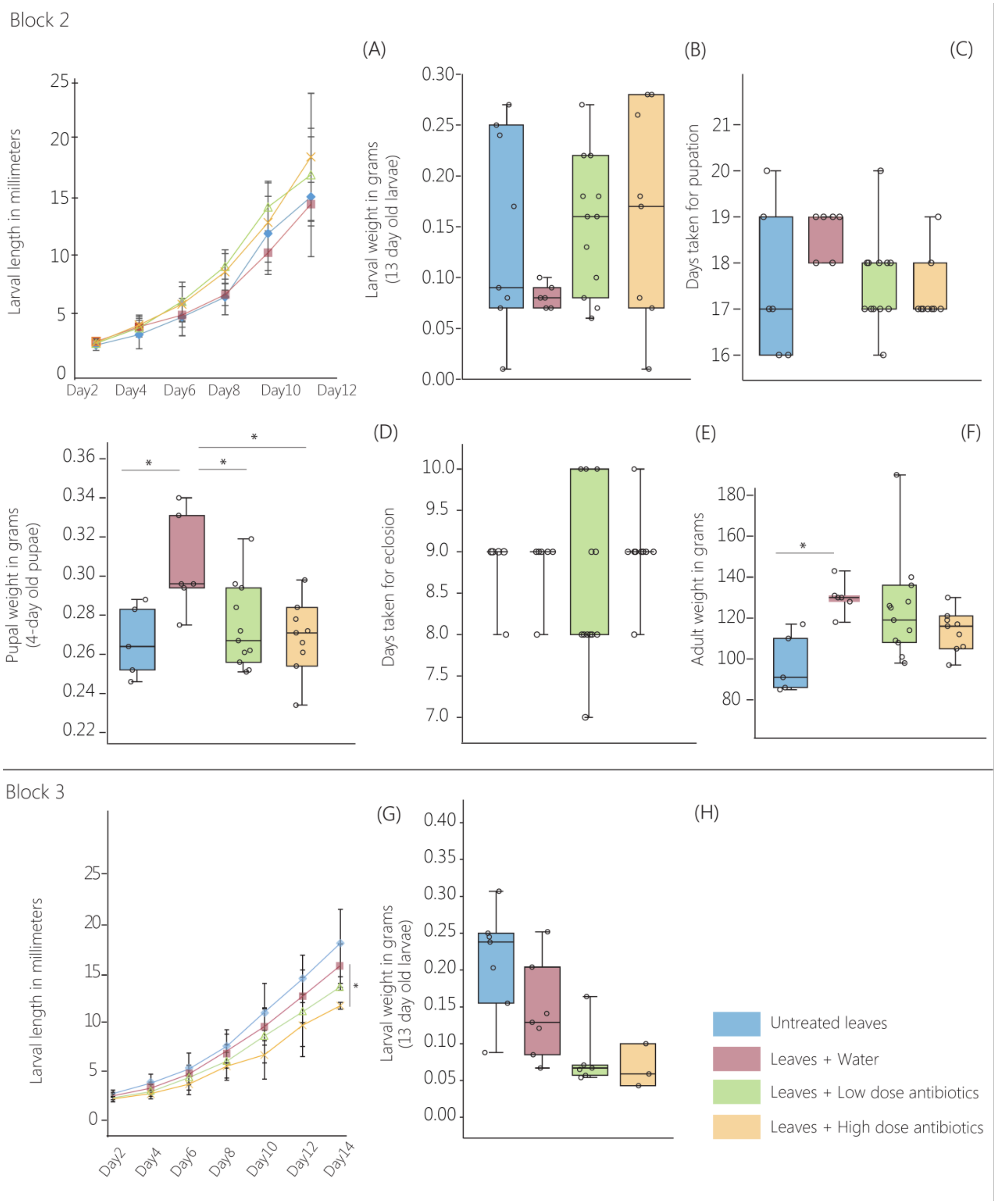

### SUPPLEMENTARY METHODS

#### Testing for bacterial contamination

As shown in earlier report [1], DNA extraction kits can introduce bacterial contamination. Hence, we tested our DNA extraction kits for possible contaminants, performing a mock DNA extraction (without any animal tissue) as a negative control. We performed 2 rounds of PCR as part of the library preparation protocol and quantified the amount of amplified PCR product using Qubit, a sensitive method for DNA quantification. For all larval samples, we obtained ~80 ng/µl DNA per sample; however we could not detect any amplification from the negative control. This suggested that the probability of bacterial contamination from our DNA extraction kits was very low.

#### Analyzing larval bacterial communities using QIIME

For obtaining bacterial OTUs, we used the following commands in QIIME:

- *multiple_join_paired_ends.py* (Join forward and reverse reads)
- *multiple_split_libraries_fastq.py* (q score >29) (Filter low quality reads)
- *identify_chimeric_seqs.py* and *filter_fasta.py* (Identify and filter chimeric sequences)
- *pick_open_reference_otus.py* (Pick bacterial OTUs)

For representing the dominant bacterial community members of *D. chrysippus* and *A.merione* larvae, we selected the five most abundant bacterial OTUs as described earlier [2] (figure 1, panels A1-C1).

#### Quantifying bacterial abundance using quantitative PCR (qPCR)

We set up a 10µl PCR reaction for each sample using 5µl SYBR green (Maxima SYBR Green/ROX qPCR Master Mix (2X), Thermo Fischer Scientific), 1µl forward and reverse primer each (10µM), 1 µl larval DNA extract (~200ng DNA) and 2µl water. We set up qPCR reactions in 384 well plates (MicroAmp™ Optical 384-Well Reaction Plate with Barcode, Applied Biosystems). We performed genomic qPCR using the ViiA™- 7 Real-Time PCR System (Applied Biosystems) as follows: 5 min at 95°C followed by 40 cycles [45 sec at 95°C, 30 sec at 60°C, 45 sec at 72°C] and recorded the Ct values (cycle threshold) for each sample. We calculated the ΔCt (internal control Ct – target gene Ct) and quantified the abundance of bacteria in each sample using the formula ($2^{\wedge\, -\Delta Ct}$), normalizing the amplification of the bacteria-specific 16S rRNA gene with that of the butterfly-specific 18S rRNA gene. We performed this normalization to compare the bacterial load per unit amount of host DNA. We used previously reported primers for qPCR [3–7]. The reverse primer for host-specific 18S rRNA gene amplification was designed by Kunte lab. The relevant primer sequences are given below.

| Target gene | Forward primer (5' – 3') | Reverse primer (5' – 3') |
|---|---|---|
| 18S rRNA gene (Host specific) | CGGCTACCACATCCAAGGAA | GGCCTCGTAAGAGTCCCGTAT |
| 16S rRNA gene (Gammaproteobacteria) | CMATGCCGCGTGTGTGAA | ACTCCCCAGGCGGTCDACYTA |
| 16S rRNA gene (Actinobacteria) | TACGGCCGCAAGGCTA | CATCCCCACCTTCCTCCG |
| 16S rRNA gene (Firmicutes) | ACCATGCACCACCTGTC | TGAAACTYAAAGGAATTGACG |

**Measuring host fitness**

To test the impact of bacterial elimination on the host development, we measured and compared different fitness proxies across control and treated groups. Before starting the manipulative experiments, we distributed approximately equal number of eggs in each treatment. However, all eggs in each group did not hatch. Thus, we ended up with a slightly different number of larvae in each treatment group (see supplementary tables). Also, the number of individuals in each treatment group varied slightly while measuring different fitness proxies for different developmental stages, for the following reasons. A few larvae died during the course of development and thus we could not measure their fitness at the pupal or adult stage. In rare cases, adults fell down from the pupal case right after eclosion, even before they could expand their wings. We were unsure if these adults could release meconium (metabolic waste) completely. To avoid overestimation of adult weight due to incomplete meconium release, we did not measure their body weight. In a few cases, individuals died in the pupal case and adult fitness could not be measured. None of these instances were treatment-specific. For instance, larval mortality did not vary significantly and consistently across treated and control groups across experimental blocks (Fisher's exact test, $p>0.05$, see table 1). Finally, we also stored ~2-3 larvae from a few experimental blocks for analyzing larval bacterial communities and thus could not measure their fitness at the pupal and adult stage. The number of individuals included in each treatment within each block is shown in supplementary table S4.

**SUPPLEMENTARY TABLES**

**Table S1- Impact of dietary sterilization on the fitness of *D. chrysippus***

Table 1.1 shows the treatments included in each experimental block. Table 1.2 shows the results of analyses of fitness measurements across treatments. Asterisks indicate significant variation between the control group (treatment A) and other treatments (Generalized linear model, Tukey's post hoc test for multiple comparisons, *p < 0.05), with the direction of the difference as indicated. "E" represents the "estimate" from the Tukey's post-hoc test and is reported only for the comparisons that are significant. Non-significant comparisons are indicated as "ns", and fitness proxies that were not determined are indicated by "nd". See table S4 for the exact number of replicates for each treatment group and fitness measurement.

**1.2: Treatments included in each experimental block**

| **Treatments** | **A**<br>**Unsterile Diet** | **B**<br>**Sterile Diet + Leaf flora** | **C**<br>**Sterile Diet + Frass flora** | **D**<br>**Sterile Diet** |
|---|---|---|---|---|
| Block 1 | ✓ | nd | nd | ✓ |
| Block 2 | ✓ | ✓ | ✓ | ✓ |
| Block 3 | ✓ | ✓ | ✓ | ✓ |

**1.1: Impact of dietary sterilization and microbial reintroduction on *D. chrysippus* fitness**

| **Experimental blocks** | **n per block** | **Larval Length** | **Larval Weight** | **Larval Span** | **Pupal Weight** | **Pupal span** | **Adult Weight** |
|---|---|---|---|---|---|---|---|
| Block 1 | 8-9 | *D<A<br>p= 9.44e-05<br>E = -9.1 | *D<A<br>p=0.0231<br>E = -0.10 | *D>A<br>p=0.025<br>E= 11.2 | ns | ns | ns |
| Block 2 | 6-13 | *D<A<br>p=0.04<br>E= 3.1 | nd | ns | ns | ns | *D<A<br>p=0.04<br>E = -0.06 |
| Block 3 | 6-8 | ns | nd | *B>A<br>p=0.001<br>E=3.6<br><br>*D>A<br>p=0.01<br>E= 2.8 | ns | ns | ns |

**Table S2- Impact of antibiotic treatment on the fitness of *D. chrysippus***

Table 2.1 shows the treatments included in each experimental block. Table 2.2 shows the results of analyses of fitness measurements across treatments (fitness of antibiotic treated groups is compared with group B; generalized linear model, Tukey's post- hoc test for multiple comparisons). Asterisks indicate significant variation (*p<0.05). "E" represents the "estimate" from the Tukey's post hoc test and is reported only for the comparisons that are significant. Replicate size per experimental block is represented as a range (n per block). Non-significant comparisons are indicated as "ns", and fitness proxies that were not determined are indicated by "nd". See table S4 for the exact number of replicates for each treatment group and fitness measurement.

**2.1: Treatments included in each experimental block**

| **Treatments** | **A** | **B** | **C** | **D** |
|---|---|---|---|---|
| | **Untreated leaves** | **Leaves + Water** | **Leaves +Low Dose antibiotic** | **Leaves + High Dose antibiotic** |
| Block 1 | ✓ | ✓ | ✓ | ✓ |
| Block 2 | ✓ | ✓ | ✓ | ✓ |
| Block 3 | ✓ | ✓ | ✓ | ✓ |
| Block 4 | ✓ | ✓ | ✓ | ✓ |

**2.2: Impact of the antibiotic treatment on *D. chrysippus* fitness**

| **Experimental blocks** | **n per block** | **Larval Length** | **Larval Weight** | **Larval Span** | **Pupal Weight** | **Pupal span** | **Adult Weight** | **Larval digestion efficiency** |
|---|---|---|---|---|---|---|---|---|
| Block 1 | 5-8 | ns | ns | ns | ns | ns | ns | nd |
| Block 2 | 4-18 | ns | ns | ns | ns | ns | ns | *B>C p= 0.04, E= -0.15<br>Weight gained by<br>7-day old larvae in 24 hrs. |
| Block 3 | 9-25 | ns | ns | ns | ns | ns | ns | ns |
| Block 4 | 4-13 | ns | ns | ns | ns | ns | ns | ns |

**Table S3- Impact of antibiotic treatment on the fitness of *A. merione***

Table 3.1 shows treatments included in each experimental block. Table 3.2 shows the results of analyses of fitness measurements across treatments (fitness of antibiotic treated groups is compared with group B, generalized linear model, Tukey's post- hoc test for multiple comparisons). Asterisks indicate significant variation (*p<0.05). "E" represents the "estimate" from the Tukey's post-hoc test and is reported only for the comparisons that are significant. Non-significant comparisons are indicated as "ns", and fitness proxies that were not determined are indicated by "nd". See table S4 for the exact number of replicates for each treatment group and fitness measurement.

**3.1: Treatments per block**

| **Treatments** | **A**<br>**Untreated leaves** | **B**<br>**Leaves + Water** | **C**<br>**Leaves + Low Dose antibiotic** | **D**<br>**Leaves + High Dose antibiotic** |
|---|---|---|---|---|
| Block 1 | ✓ | ✓ | ✓ | ✓ |
| Block 2 | ✓ | ✓ | ✓ | ✓ |
| Block 3 | ✓ | ✓ | ✓ | ✓ |

**3.2: Impact of the antibiotic treatment on *A. merione* fitness**

| **Experimental blocks** | **n per block** | **Larval Length** | **Larval Weight** | **Larval Span** | **Pupal Weight** | **Pupal span** | **Adult Weight** |
|---|---|---|---|---|---|---|---|
| Block 1 | 9-16 | ns | ns | ns | ns | ns | ns |
| Block 2 | 5-12 | ns | ns | ns | B>D* p=0.005, E= -0.04<br>B>C* p=0.02, E= -0.03<br>B>A* p=0.01, E= 0.04 | ns | B>A* p=0.01, E= 32 |
| Block 3 | 3-8 | B>D* p=0.04, E= -4.1 | ns | nd | nd | nd | nd |

**Table S4- Replicate sizes across different experimental blocks**

Tables 4.1, 4.2 and 4.3 show the number of larvae tested in each experimental block. Fitness proxies that were not determined are represented as “nd”. See supplementary tables S1–S3 for a description of treatments.

**4.1 Testing the impact of dietary sterilization on the fitness of *D. chrysippus***

| | Block 1 | | Block 2 | | | | Block 3 | | | |
|---|---|---|---|---|---|---|---|---|---|---|
| **Treatments** | **A** | **D** | **A** | **B** | **C** | **D** | **A** | **B** | **C** | **D** |
| Number of larvae at the beginning of the experiment | 9 | 9 | 13 | 13 | 12 | 15 | 11 | 11 | 11 | 13 |
| Larval Length | 9 | 9 | 13 | 13 | 12 | 15 | 10 | 11 | 11 | 12 |
| Larval Weight | 9 | 9 | nd | | | | nd | | | |
| Larval Span | 8 | 9 | 10 | 7 | 9 | 12 | 6 | 6 | 8 | 7 |
| Pupal Weight | 9 | 9 | 10 | 7 | 7 | 7 | 6 | 5 | 7 | 5 |
| Pupal Span | 9 | 9 | 9 | 6 | 9 | 10 | 6 | 6 | 8 | 7 |
| Adult Weight | 9 | 8 | 10 | 6 | 7 | 7 | 6 | 6 | 7 | 7 |

**4.2 Testing the impact of antibiotic treatment on the fitness of *D. chrysippus***

| | Block 1 | | | | Block 2 | | | | Block 3 | | | | Block 4 | | | |
|---|---|---|---|---|---|---|---|---|---|---|---|---|---|---|---|---|
| **Treatments** | **A** | **B** | **C** | **D** | **A** | **B** | **C** | **D** | **A** | **B** | **C** | **D** | **A** | **B** | **C** | **D** |
| Number of larvae at the beginning of the experiment | 5 | 5 | 8 | 8 | 14 | 16 | 15 | 18 | 29 | 26 | 23 | 25 | 13 | 13 | 11 | 13 |
| Larval Length | 5 | 5 | 8 | 8 | 13 | 15 | 14 | 18 | 24 | 25 | 16 | 19 | 13 | 13 | 8 | 12 |
| Larval Weight | 5 | 5 | 8 | 8 | 9 | 15 | 14 | 16 | 19 | 17 | 15 | 18 | 13 | 13 | 7 | 12 |
| Larval Span | 5 | 5 | 8 | 8 | 12 | 15 | 14 | 18 | 23 | 17 | 16 | 15 | 13 | 13 | 8 | 12 |
| Larval digestion efficiency | nd | | | | 8 | 13 | 14 | 15 | 20 | 14 | 15 | 15 | 8 | 6 | 4 | 8 |
| Pupal Weight | nd | | | | 6 | 8 | 11 | 9 | 22 | 19 | 20 | 21 | 11 | 11 | 8 | 11 |
| Pupal Span | 5 | 5 | 8 | 8 | 12 | 15 | 14 | 18 | 22 | 19 | 19 | 19 | 13 | 13 | 6 | 10 |
| Adult Weight | 5 | 5 | 8 | 8 | 12 | 14 | 12 | 18 | 20 | 15 | 9 | 13 | 11 | 12 | 4 | 9 |

**4.3 Testing the impact of antibiotic treatment on the fitness of *A. merione***

| | Block 1 | | | | Block 2 | | | | Block 3 | | | |
|---|---|---|---|---|---|---|---|---|---|---|---|---|
| **Treatments** | **A** | **B** | **C** | **D** | **A** | **B** | **C** | **D** | **A** | **B** | **C** | **D** |
| Number of larvae at the beginning of the experiment | 18 | 17 | 18 | 20 | 11 | 6 | 12 | 11 | 8 | 8 | 9 | 10 |
| Larval Length | 16 | 14 | 14 | 14 | 7 | 6 | 12 | 11 | 8 | 7 | 6 | 3 |
| Larval Weight | 16 | 13 | 12 | 13 | 8 | 6 | 12 | 8 | 8 | 7 | 6 | 3 |
| Larval Span | 13 | 10 | 9 | 11 | 6 | 6 | 12 | 9 | nd | | | |
| Pupal Weight | 13 | 10 | 9 | 11 | 5 | 6 | 11 | 9 | nd | | | |
| Pupal Span | 13 | 10 | 10 | 11 | 5 | 6 | 12 | 9 | nd | | | |
| Adult Weight | 13 | 9 | 9 | 11 | 5 | 6 | 12 | 9 | nd | | | |
